# Generating three-dimensional genome structures with a variational quantum algorithm

**DOI:** 10.1101/2025.04.06.647452

**Authors:** Andrew Jordan Siciliano, Zheng Wang

## Abstract

Chromosome conformation capture (3C) experiments have revealed the underlying spatial interactions that govern three-dimensional (3D) genome organization and topology. Detecting 3D contacts between genomic loci considerably enhances our understanding of fundamental regulatory processes. Modeling 3D structures from experimental contact matrices can further contextualize the relationship between 3D genome organization and regulation. While classical algorithms have been successful in reconstructing genomic conformations, we investigate the prospect of quantum computation to aid in modeling the conformational space. In this context, we propose a novel variational quantum algorithm to model the distribution of 3D genomic structures from experimental contact data. Through rigorous evaluations, we demonstrate the capability of our algorithm to sample ensembles of viable 3D conformations that agree well with experimental and simulated contact data. Furthermore, we extend our methodology to model the conformational space of a single cell or a population of cells. In the advent of sufficient quantum utility, the insights gained from this study can serve as a foundation for investigating high-resolution, large-scale ensembles of genomic conformations through generative variational quantum algorithms.

## 1 Introduction

Chromosome conformation capture (3C) experiments detect topological and organizational properties of the entire physical genome^1^, and captured contacts relate directly to the spatial proximity between pairs of genomic loci. More recently developed experimental protocols, such as the Hi-C^2,3^ and Micro-C^4^ experiments have provided the underlying spatial organization of entire genomes with unprecedented details. Single-Cell Hi-C^5^ is an extension of the Hi-C experiment that provides us with contacts at the level of individual cells. Single-cell genomic conformations have a high degree of variability^5^, with respective contacts providing a deeper understanding of cell-cycle dynamics^6^ and genome regulation processes^7^. 3C experiments and their derivatives reveal structural features of chromatin organization, such as topological domains^8^ and chromatin loops^9^. These features have proven invaluable for understanding promoter-promoter and promoter-enhancer interactions, gene activation processes, gene expression, transcriptional regulation, and disease development and progression^8–27^.

Captured genomic contacts are inversely correlated with pairwise three-dimensional (3D) target distances. We can use inverse exponential functions, such as 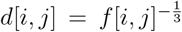 with *d*[*i, j*] being the target distance and *f*[*i, j*] being the contact frequency captured between loci *i* and *j*, to produce target distance matrices (distograms) of the full genomic structures^28–30^. However, to understand the relations between genomic structure and function, modeling (simulating) the structure of 3D genomes is necessary to gain further insights^31^. For example, structural models can help decipher the relationship between 3D conformations and gene regulation, gene expression, and enhancer activities^31–36^. Computational methods have been successful in approximately reconstructing the spatial locations of 3D genomic structures from the captured contacts (or target distances), with common approaches including numerical optimization and simulated annealing^27–30,36–43^.

Many 3D conformations can fit the experimentally determined contacts, thus producing ensembles of viable genomic conformations is of high interest. Ensembles are commonly generated via parallel optimization routines, such as in deconvolution methods^35,41^, or repeated runs of a proposed algorithm from different starting points or random seeds, such as in sampling methods^38,44^. Probabilistic modeling has also been shown effective in producing diverse structural ensembles^41,42,44^. However, many approaches become computationally expensive due to lengthy simulation (optimization) times or large memory requirements. Oftentimes generating even moderately sized ensembles can demand significant resource utilization, especially so for high-resolution contact data.

In the disciplines of bioinformatics and computational biology, quantum computing has, in recent times, begun to be explored^45–49^. Many proposed algorithms offer promising utility. For example, quantum algorithms have been proposed for de novo genome assembly^50–52^, mRNA codon optimization^53^, the RNA folding problem^54,55^, and protein binding site identification^56^. Notably, an adaption of the Variational Quantum Eigensolver^57^ that utilizes Conditional Value-at-Risk (CVaR)^58^ has been applied to the protein folding problem. The model proposed by Ref.^59^ was able to successfully fold small peptides on a tetrahedral lattice, with their proposed algorithm achieving results comparable to those obtained through x-ray crystallography^60^. However, the feasibility of tackling the protein folding problem on near-term devices is still an open problem with potential barriers being explored^61,62^.

Variational quantum algorithms (VQA) are heuristic-based algorithms heavily researched for quantum chemistry and combinatorial optimization. VQAs involve both classical and quantum computation, linked in a feedback loop. An objective function ℒ is computed over a set of sampled measurements, and parameterized circuits (ansatz) are variationally adjusted to optimize ℒ. The applications of VQAs on Noisy Intermediate Scale Quantum (NISQ)^63^ devices are suggested to be promising, due to shallow circuit depth requirements and some resilience to noise^64^. However, issues of barren plateaus^65,66^ and objective function landscape difficulties^67,68^ are intrinsic problems associated with training VQAs. It is still unclear whether advantages in the near term are feasible^69^, however, potential mitigation strategies are being explored with some success^70–78^. Even with the aforementioned obstacles, given improvements in hardware and error correction technologies, the prospective real-world usefulness of VQAs could be wide-reaching^79,80^.

The Variational Quantum Eigensolver^57^ (VQE) is a fundamental VQA, which attempts to approximate the ground (lowest energy) state of a given Hamiltonian ℋ. VQE can be used for combinatorial optimization problems when is ℋ equivalent to a pseudo-boolean polynomial^81^. A parameterized quantum state is prepared using an Ansatz circuit, with parameters variationally adjusted to minimize the expectation of the energy of the system. Typically, ℋ is decomposed into a set of Pauli strings, and the expected energy is computed using the Hamiltonian averaging technique^57,82^. The procedure for a wide variety of VQAs is similar to that of VQE, with the difference being how the objective function is defined and computed. While the goal of VQE is to approximate the ground state of the Hamiltonian, sometimes it is preferred to model a specific target distribution, such as in quantum generative modeling. We can define an objective function whose minimum guides the quantum state toward the target distribution of interest. Objective functions need not be fully classical and can also be designed to compare sample data points using kernel and discrepancy methods^83^.

Quantum machine learning (QML)^84^ has applications for both supervised and unsupervised learning. It is natural to employ quantum systems as generative models, as they are fundamentally probabilistic. Quantum generative models (QGM) are VQAs that aim to embed a target distribution in the quantum system using parameterized circuits. It has been shown that under certain conditions QGMs can provide clear benefits with respect to expressibility and sampling efficiency^85–87^. Determining the long-term viability of VQAs in the context of generative modeling is still an open problem, however, current research has suggested promising insights and perspectives on the future applications of QGMs^83,88–90^.

We developed a variational quantum algorithm that aims to model the conformational space of 3D genomic structures. To the best of our knowledge, this is the first time quantum computing has been proposed for the 3D genome reconstruction problem. We implemented and investigated a mathematical model specifically designed to simulate high-likelihood ensembles of genomic conformations from bulk and single-cell Hi-C data on a quantum computer. In the advent of sufficient quantum utility^63,91^, our proposed algorithm could offer further insights into the space of viable 3D genomic conformations, advancing our understanding of 3D genomic organization and its role in various biological processes.

## 2 Methods

### 2.1 Problem statement and objective function

We formulate the 3D genome reconstruction problem in terms of contacts captured from a 3C or derivative experimental protocol. We assume Hi-C as the experimental protocol of choice, and the aim is to find *N* three-dimensional (3D) coordinates whose pairwise Euclidean distances correlate well with the contact frequency matrix *f* ∈ ℝ^*N ×N*^. We can simplify the problem further by assuming contacts are binary^38^, where the positive label could be thought of as a detected chromatin loop^9^.

Genomic structures are threaded through a cubic lattice, where each bead *i* can move to the 8 possible corners of a cube centered at bead *i* − 1, see Figure 1. We map genomic structures of length *N* to the state of 3(*N* −2) qubits, with the first two beads 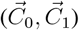 fixed, and the distance between beads *i*−1 and *i* (unit length) as 1. We define the coordinate 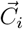 (see Equations 1 and 2) as a unit vector composed of three binary (spin) variables 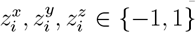,corresponding to the 8 corners of a cube, translated by 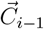,see Figure 1.

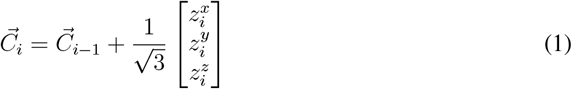

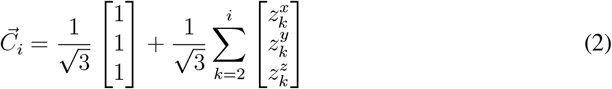

**Figure 1:**
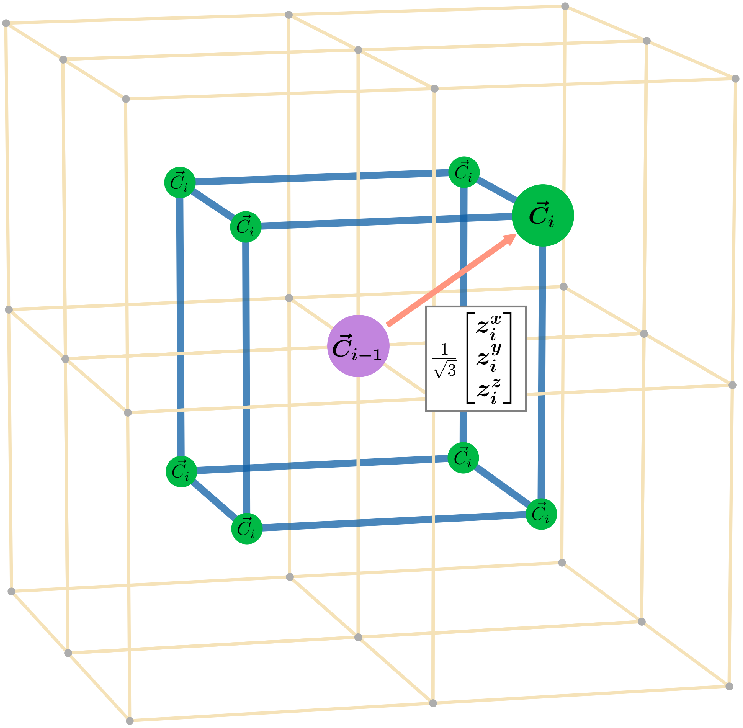
The eight possible locations for *C*_*i*_ (green), with respect to the cube centered on *C*_*i*−1_ (purple).

The squared Euclidean distance is defined as *E*[*i, j*], see Equation 3. Note that by the properties of the lattice *E*[*i, j*] is bounded by (*i* − *j*)^2^, i.e. *E*[*i, j*] <= (*i* − *j*)^2^.

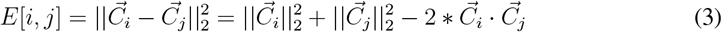

From the Hi-C experiment we can construct a contact matrix *π*_*c*_. A contact implies that the distance between the beads is less than or equal to some pre-defined radius *r*:

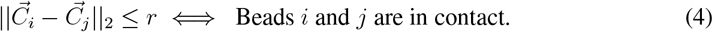

Let us define the entries of the contact matrix as *π*_*c*_[*i, j*], which gives the probability of a contact between beads *i* and *j*. As previously mentioned, for simplicity we assume *π*_*c*_[*i, j*] ∈ {0, 1}, making *π*_*c*_ a binary matrix. We construct the likelihood of *E*[*i, j*] given *π*_*c*_[*i, j*] (Equation 7) in terms of Equations 5 and 6, which correspond to attractive (*E*[*i, j*] should approach *r*^2^) and neutral propensities, respectively.

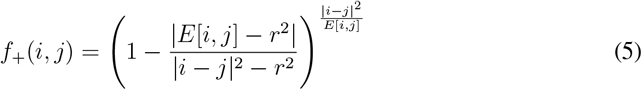

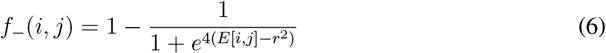

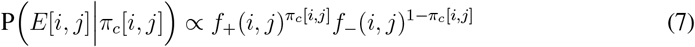

When a contact is not detected by the Hi-C experiment, i.e. *π*_*c*_[*i, j*] = 0, the likelihood is an S-curve (*f*_−_) that gives approximately equal plausibility for all distances except those that approach or surpass the contact threshold *r*. When *π*_*c*_[*i, j*] = 1, the likelihood is higher for distances near the contact threshold *r* and lower otherwise (*f*_+_), see Figure 2.

**Figure 2:**
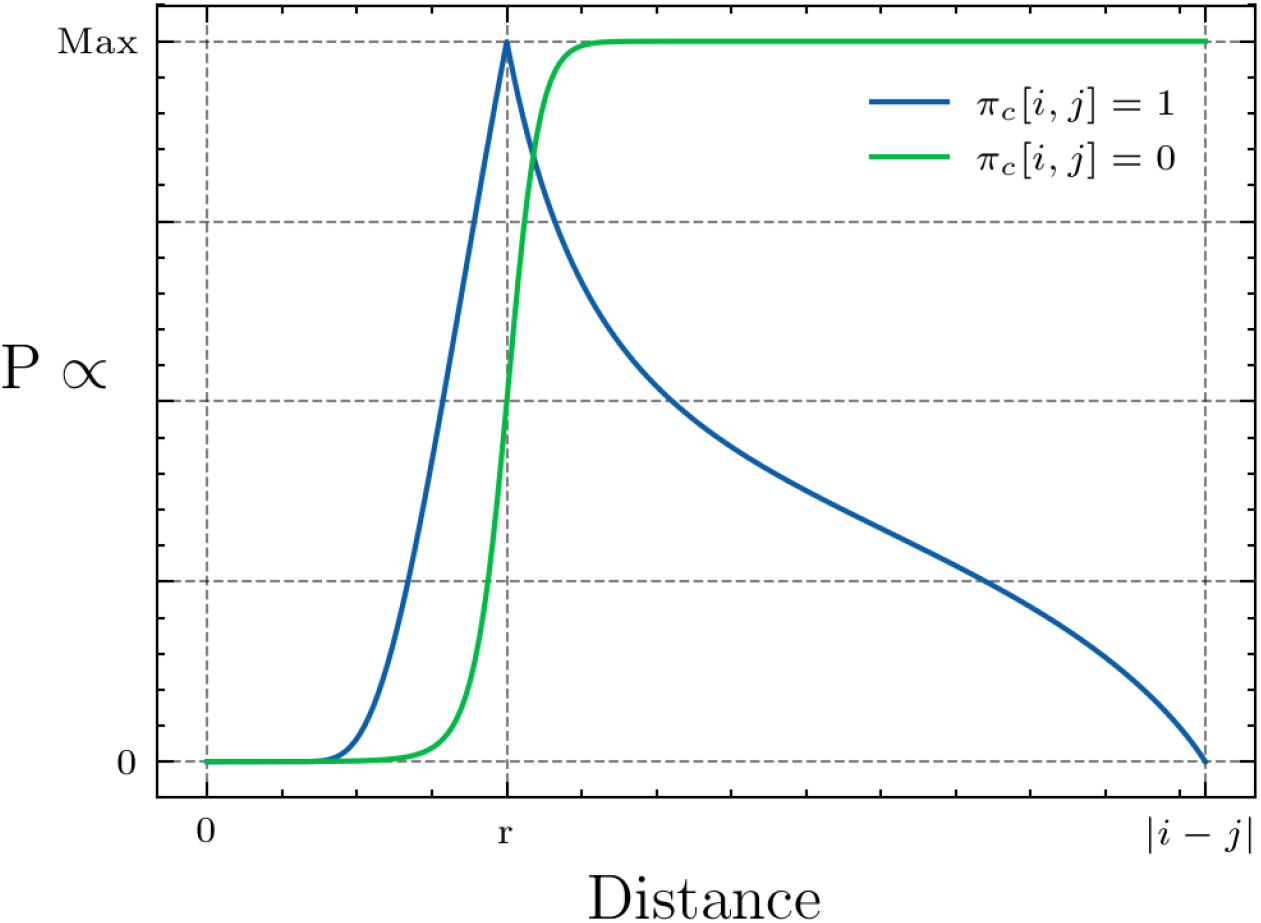
The likelihood of the distance 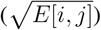 between beads *i* and *j* given *π*_*c*_[*i, j*].

For a given squared distance matrix *E*_*k*_ associated with a structure 𝕊_*k*_ ∈ ℝ^*N ×*3^, we model the likelihood of 𝕊_*k*_ as:

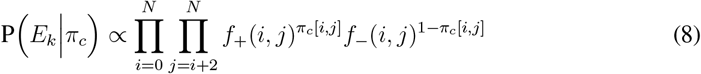

The log-likelihood of Equation 8 is defined as the following:

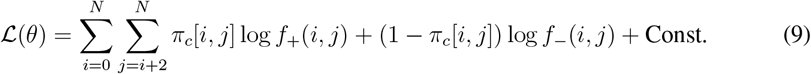

We further expand our model to account for the case of physical aggregations. Physical aggregations in the Hi-C experiment are the consensus non-single-cell contacts captured between genomic loci. Bulk Hi-C can be viewed as the average of single-cell Hi-C data^5^, thus Equation 9 assumes zero aggregation. To incorporate aggregation into our model, we consider the case where multiple structures are sampled at a time (per-shot measurements from the variational quantum algorithm). Equations 10 and 11 define the criteria for aggregated and non-aggregated contacts over a collection of sampled structures, respectively.

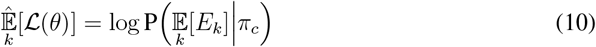

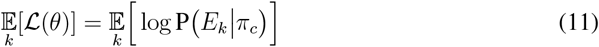

The criteria of 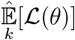 evaluates the log-likelihood of the expected squared distance matrices (aggregated distograms), as opposed to the expected log-likelihood of each individual square distance matrix, 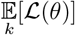. Optimizing over an objective function of the form 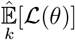 is highly applicable to variational quantum algorithms, as such functions require access to multiple samples per optimization step, aligning well with the VQA procedure. We define the generalized objective function over the sampled structures as:

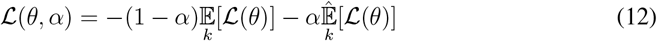

The value of *α* ∈ [0, 1] can be set depending on the level of aggregation one would like to model. Higher and lower values of *α* closer resemble the bulk and single-cell Hi-C experiments, respectively. When we increase the value of *α*, we observe an increase in the diversity and variety of sampled structures, aligning well with the relation of *α* to bulk Hi-C, for more details see Section 3. When *α* = 1, we can interpret the sampled structures as an approximate subset of single-cell structures from which the bulk Hi-C experiment was performed, i.e. a de-convolutional method. More details pertaining to the impact of *α* on the objective function landscape, learned distributions, and respective results can be found in Section 3.

Note that our model was intended for shorter-length structures (*N* < 150) due to the limitations of classical simulation and currently available quantum devices. Longer lengths could benefit from a modification to the power 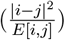 in Equation 5 to better penalize spatial clashes.

### 2.2 Choice of ansatz and optimizer

The Hamiltonian with expectation equivalent to 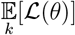 can be described entirely by Pauli Z (*σ*_*z*_) operators^92,93^. Similarly, the Hamiltonian with expectation equivalent to 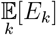 is an Ising model, and is also composed entirely of *σ*_*z*_ operators. From these observations, we can restrict our quantum state space to contain strictly real amplitudes. For simplicity, we utilized two repetitions of the RealAmplitudes ansatz provided by Qiskit^94^, see Figure 3. The number of parameters, *p*, to model *N* beads on a string is *p* = 9(*N* − 2). For the choice of optimizer, we used COBYLA^95^ since the values of *p* experimented with were relatively small. Parameters were initialized randomly over the uniform distribution with range [0, 2*π*).

**Figure 3:**
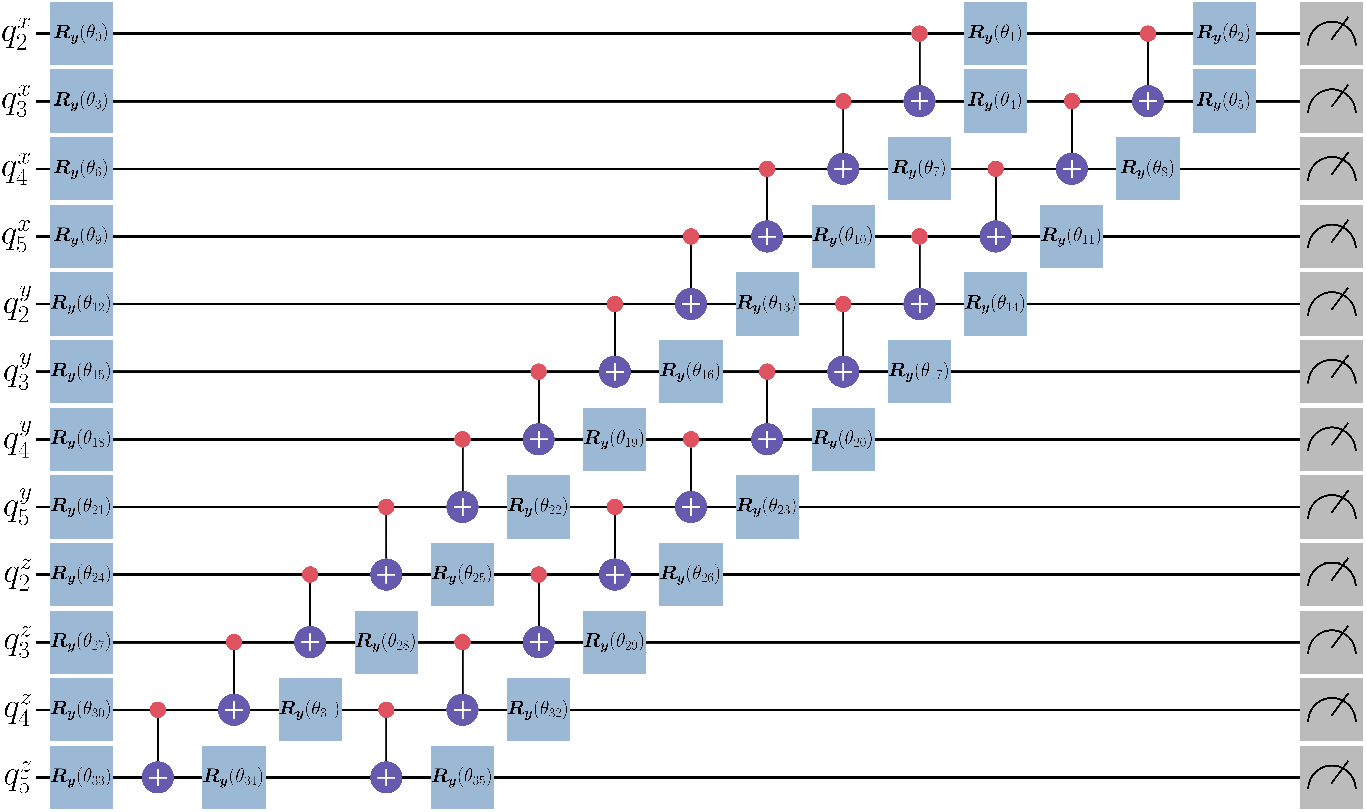
Two repititons of the RealAmplitudes Ansatz with reverse linear entanglement for a 12 qubit system (length *N* = 6 structures).

As more capable quantum machines come to fruition, it will be necessary to more thoroughly investigate both the choice of classical optimization algorithm, Ansatz, and parameter initialization strategies. Exploring these areas are outside the scope of this research. For more information on these topics and their implications with regard to variational quantum algorithms, we refer to Refs.^79,96–102^.

### 2.3 Variational quantum algorithm for generating genomic conformations

We now state the proposed variational quantum algorithm for the 3D genome reconstruction problem. First, we randomly initialize *θ* ∈ [0, 2*π*) and prepare the quantum state |*ψ*(*θ*) ⟩ using the parameterized Ansatz. Then, we sample *ρ* measurements, 𝕄, from |*ψ*(*θ*) ⟩. We map the measurements 𝕄 to structures 𝕊^*′*^ and compute the cost ℒ (*θ, α*) over 𝕊^*′*^. The cost and current *θ* are passed to a classical optimization algorithm to produce *θ*^*′*^ and update *θ* ← *θ*^*′*^. The process repeats until convergence. We can generate an ensemble of viable genomic conformations by preparing and sampling from |*ψ*(*θ*^*^)⟩. See Figure 4 and Algorithm 1.

**Figure 4:**
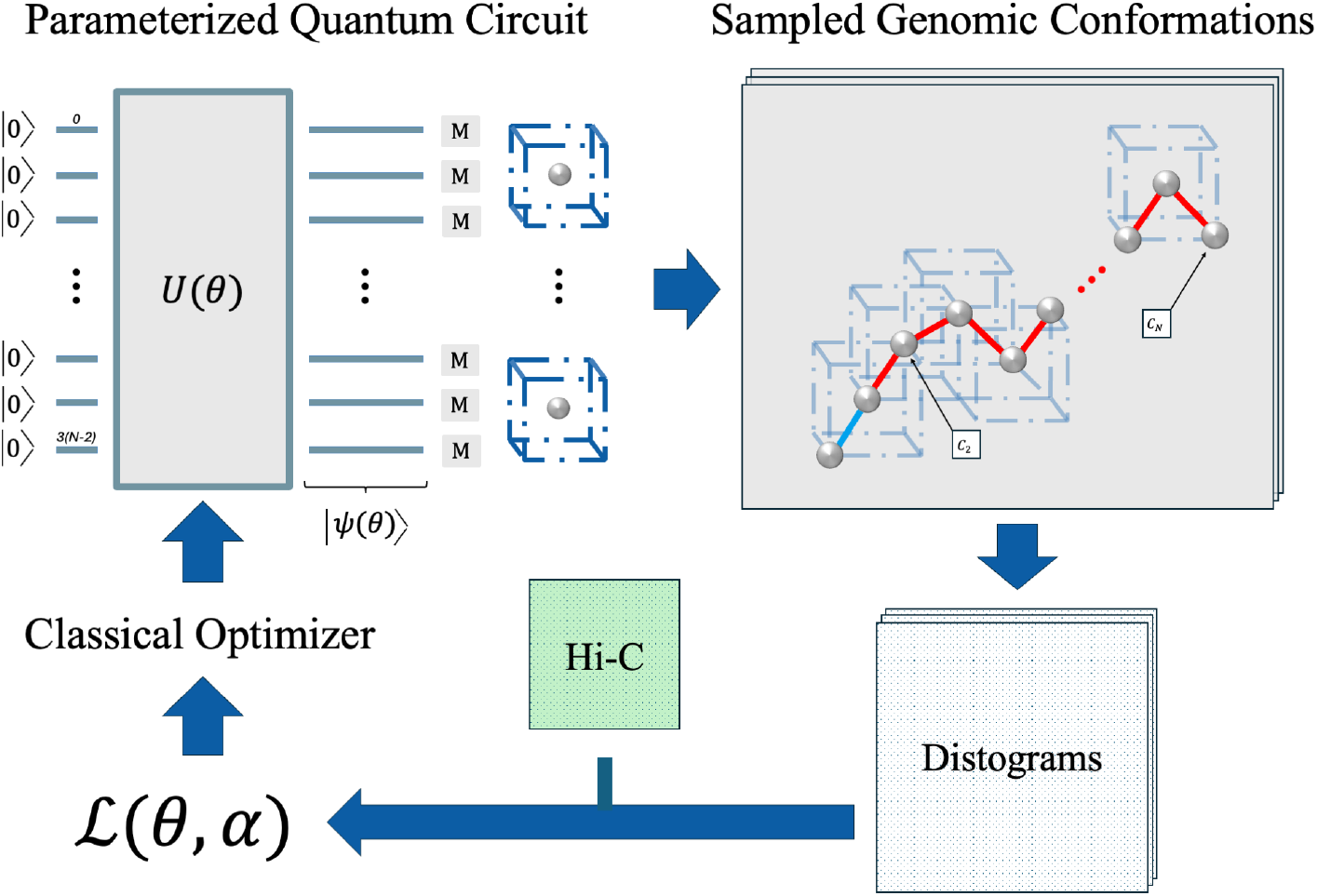
Variational quantum algorithm for generating genomic conformations.

Note that all of the Pauli strings needed to compute ℒ (*θ, α*) mutually commute with one another, thereby allowing for simultaneous measurements^103,104^. It takes *𝒪* (*N* ^2^*ρ*) time, with *ρ* being the number of measurements (samples), to compute ℒ (*θ, α*).

#### Algorithm 1 Variational Quantum Algorithm

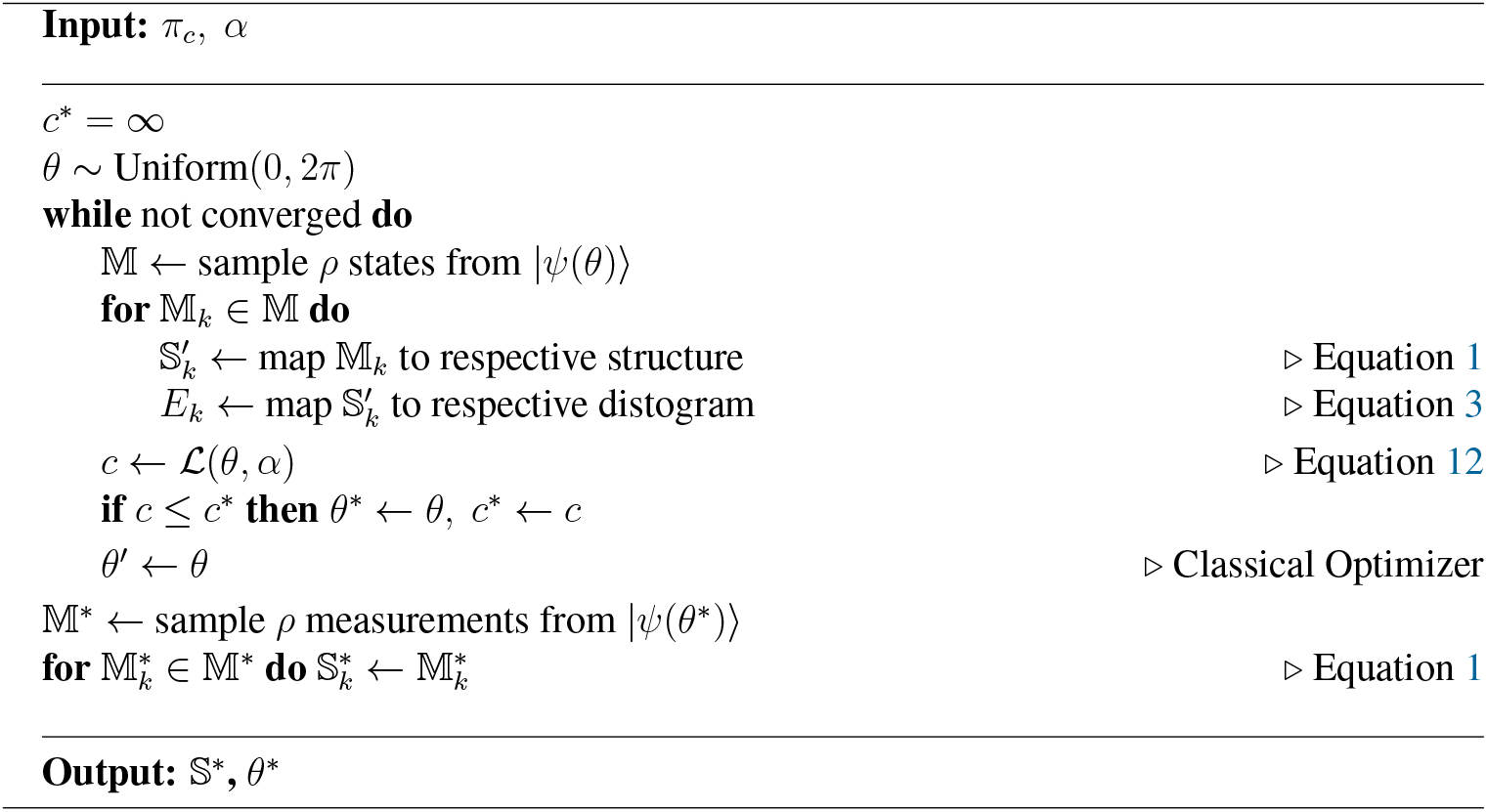

## 3 Results

### 3.1 Simulated genomic contact dataset

To evaluate whether our proposed algorithm can effectively learn the conformational space, we generated multiple simulated Hi-C datasets. Let us define the set of length *N* structures as 𝕊, with 𝕊_*k*_ ∈ ℝ^*N ×*3^ and| 𝕊 |= 2^3(*N*−2)^ = 8^*N*−2^. We were able to generate, via brute force, all possible structures with *N* ∈ [6, 11) that are free from spatial clashes (all pair-wise distances are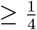). We partitioned each 𝕊 into a variety of subsets (groups) 𝒢 ⊂ 𝕊. Topological properties are extracted from each 𝒢 and used to define the ground truth for the objective function ℒ (*θ, α*).

Specifically, we defined subsets of structures through common pair-wise contacts. Two beads *i* and *j* are in contact when their Euclidean distance is within a pre-defined radius *r*, see Equation 4. For our experiments, we set *r* = 1.5. Groups are formally defined as the following:

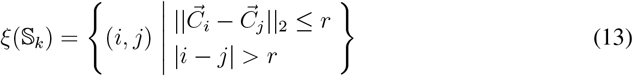

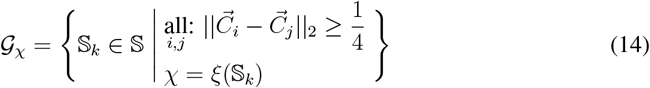

The contacts related to each group can be viewed as the result of a Hi-C experiment, where a contact implies a loop between genomic loci *i* and *j*. For a group 𝒢_*χ*_, the respective ground truth is defined as the following:

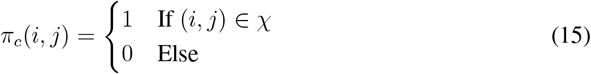

#### 3.1.1 Evaluation from simulation of quantum circuits on a classical computer

In our experiments, we set the number of shots *ρ* = 4096. We then randomly sampled targets 𝒢_*T*_ ⊂ 𝕊 for structures of lengths *N* ∈ [6, 11). We ran our VQA ten times per-target over as many targets that were computationally feasible. To verify the performance of our algorithm, and the subsequent impact of the parameter *α*, we computed multiple evaluation metrics over sampled ensembles of 4096 structures, 𝕊_*k*_ ∈ 𝕊^*^, from |*ψ*(*θ*^*^)⟩.

To directly evaluate the ability of our algorithm for finding likely conformations, for each target group we computed all of the possible theoretical likelihoods (Equation 8) across the 8^*N*−2^ states, each of which is associated with an energy level. Higher likelihoods are associated with lower energy levels and vise versa. We define the ground truth distribution tensor, *ϕ*(*π*_*c*_), as the following:

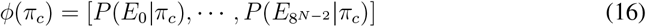

From *ϕ*(*π*_*c*_), we computed the structures in the upper *q* quantile of the energy levels. We then compared those against the 4096 sampled structures 𝕊_*k*_ ∈ 𝕊^*^. Coverage is defined as the percent of the total states in the upper *q*’th quantile, 𝕊[*ϕ*(*π*_*c*_) ≥ *Q*(*ϕ, q*)], sampled by the algorithm:

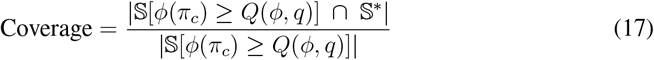

The empirical probability given *q* is the observed chance for a sampled structure 𝕊_*k*_ ∈ 𝕊^*^ to be at or above the upper quantile *q*. In Figure 5, we plot coverage and the empirical probability for quantiles in the range of [0.5, 1) for structures of length *N* = 10.

**Figure 5:**
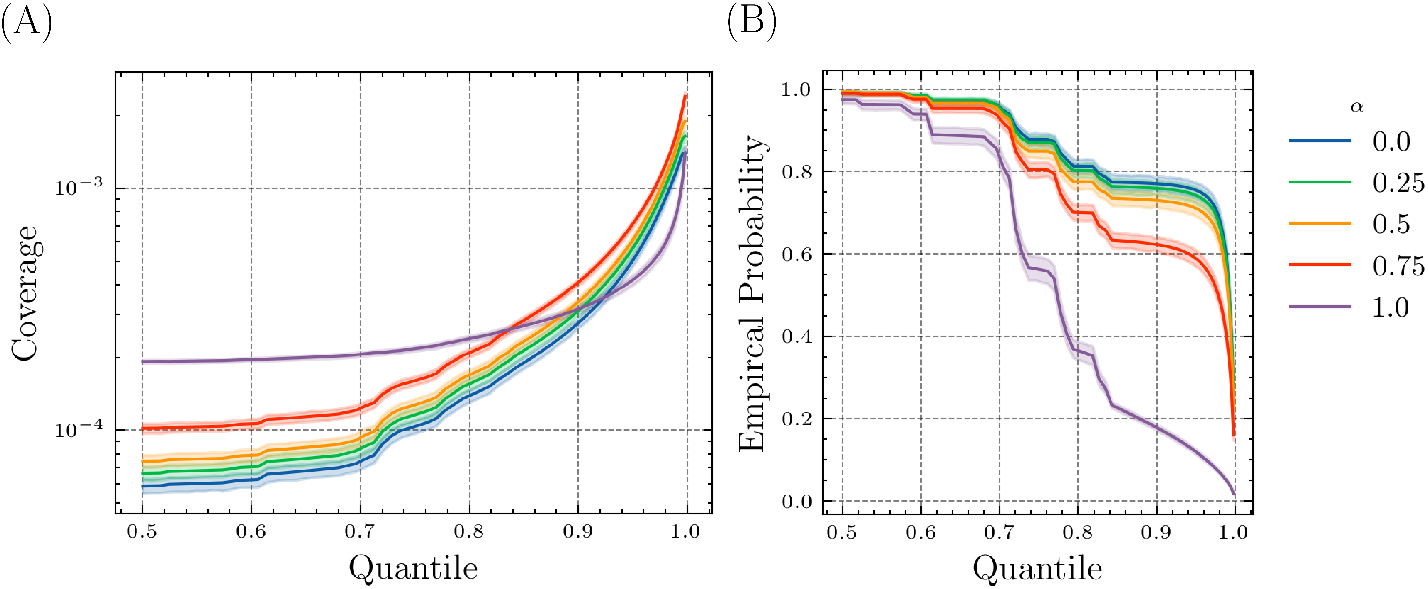
Line plot and confidence interval of the Coverage (A) and Empirical Probability (B) given *q* (quantile) for structures of length 10 over 48 target groups.

It is clear from Subfigure 5.B that the empirical likelihood of sampling a structure in the top-50th percentile (quantile) is nearly 100%. However, this chance decreases as the quantile increases for all values of *α*, with the rate of decrease being more significant for larger values of *α*. Both Subfigures 5.A and 5.B indicate that higher values of *α* can hinder the ability to sample structures with the exact target contacts, but can improve the ability sample a variety of structures with reasonable similarity to the target contacts (Bulk Hi-C).

In Figure 6 we plot the likelihood ratio 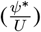 of the learned quantum distribution (*ψ*^*^) over the uniform distribution (*U*) partitioned by excitation level–equivalent to the likelihood (Equation 8). When 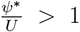,the learned probability (*ψ*^*^) of sampling a structure with a given excitation level is 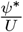 times more likely compared to random chance (*U*).

**Figure 6:**
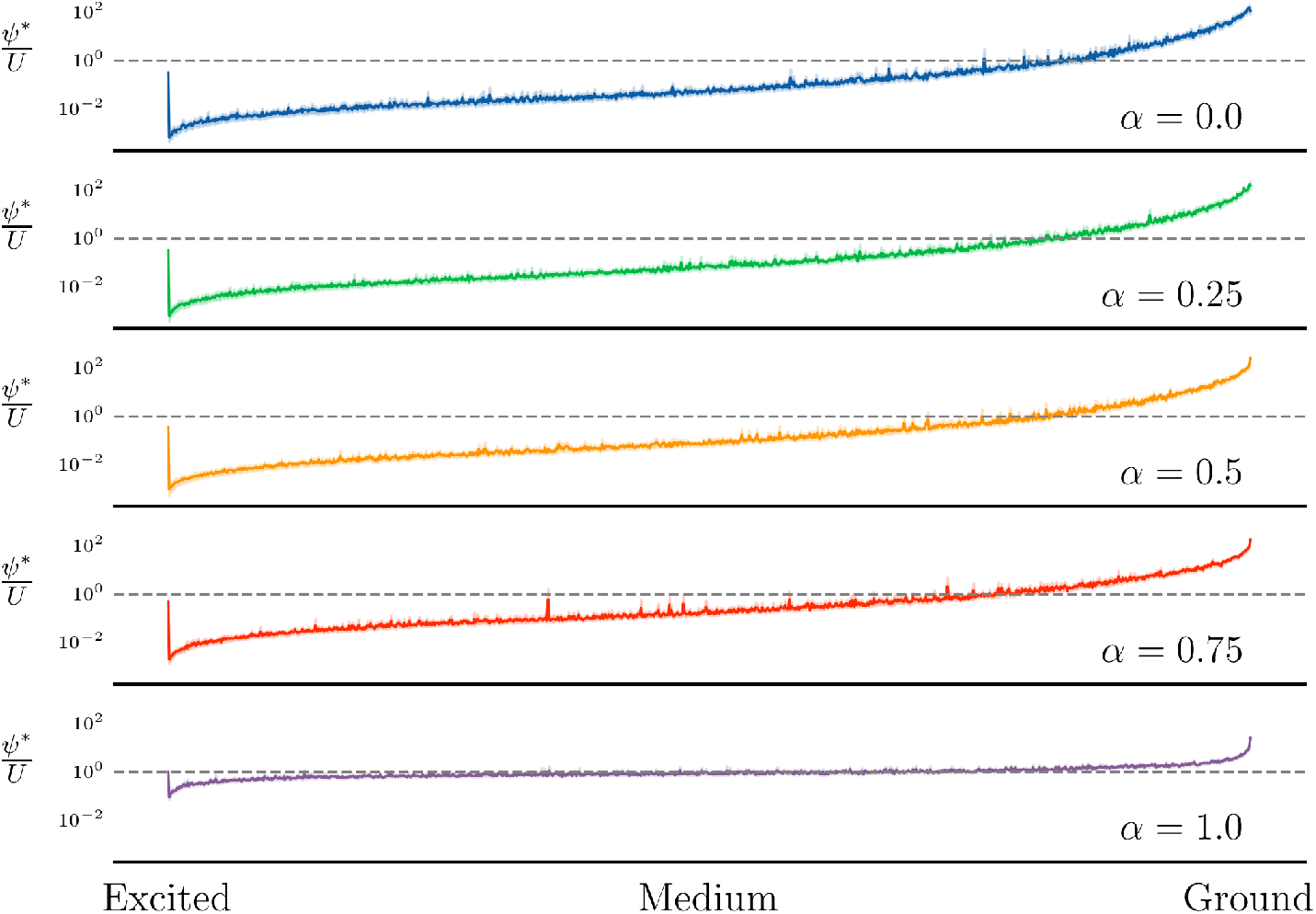
Binned line plot with confidence interval for the likelihood ratio of the learned quantum distribution over the uniform distribution, partitioned by excitation (energy) level, for length *N* = 10 structures.

What we observe is that smaller values of *α* tend to more drastically reward and penalize lower- and higher-energy solutions, respectively. Large values of *α* still penalize high-energy solutions, but are more lenient towards medium to lower-energy solutions. In other words, higher values of *α* give leeway to solutions that are not perfect, but are still reasonably likely given the contact data. This suggests that by increasing *α* we focus the model on sampling a more diverse, evenly distributed set of states across the medium to lower-energy spectrum (bulk Hi-C). Hence, as we decrease *α* we notice the sampled structures are more highly concentrated and focused around the lowest energy states of the system (single-cell Hi-C). It is worth noting that in general a large proportion of states are high-energy, see Figure 7.

**Figure 7:**
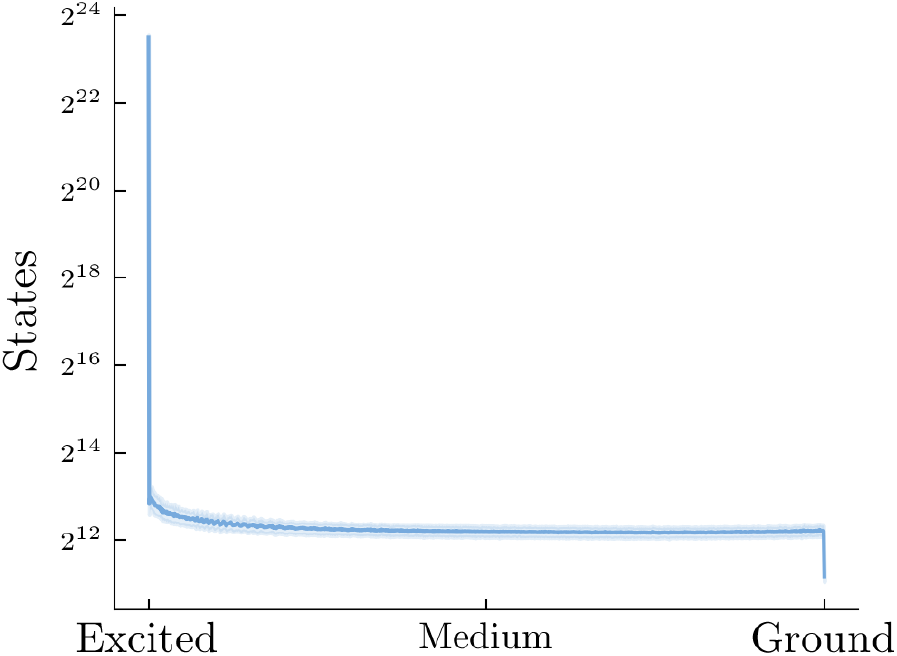
Line plot and confidence interval of the energy distribution of states, partitioned by excitation level, over 48 target groups with length *N* = 10 structures.

To further support these findings, we plot the Shannon-Entropy (*H*)^105^ curves of our VQA for structures of length *N* = 6, see Figure 8. What we notice is that as more optimization steps (lower costs) are performed, the decrease in Shannon-Entropy of the learned probability distribution is much larger for smaller values of *α*. The distribution with maximal entropy is the uniform distribution, which suggests larger values of *α* associate with more evenly distributed learned quantum states.

**Figure 8:**
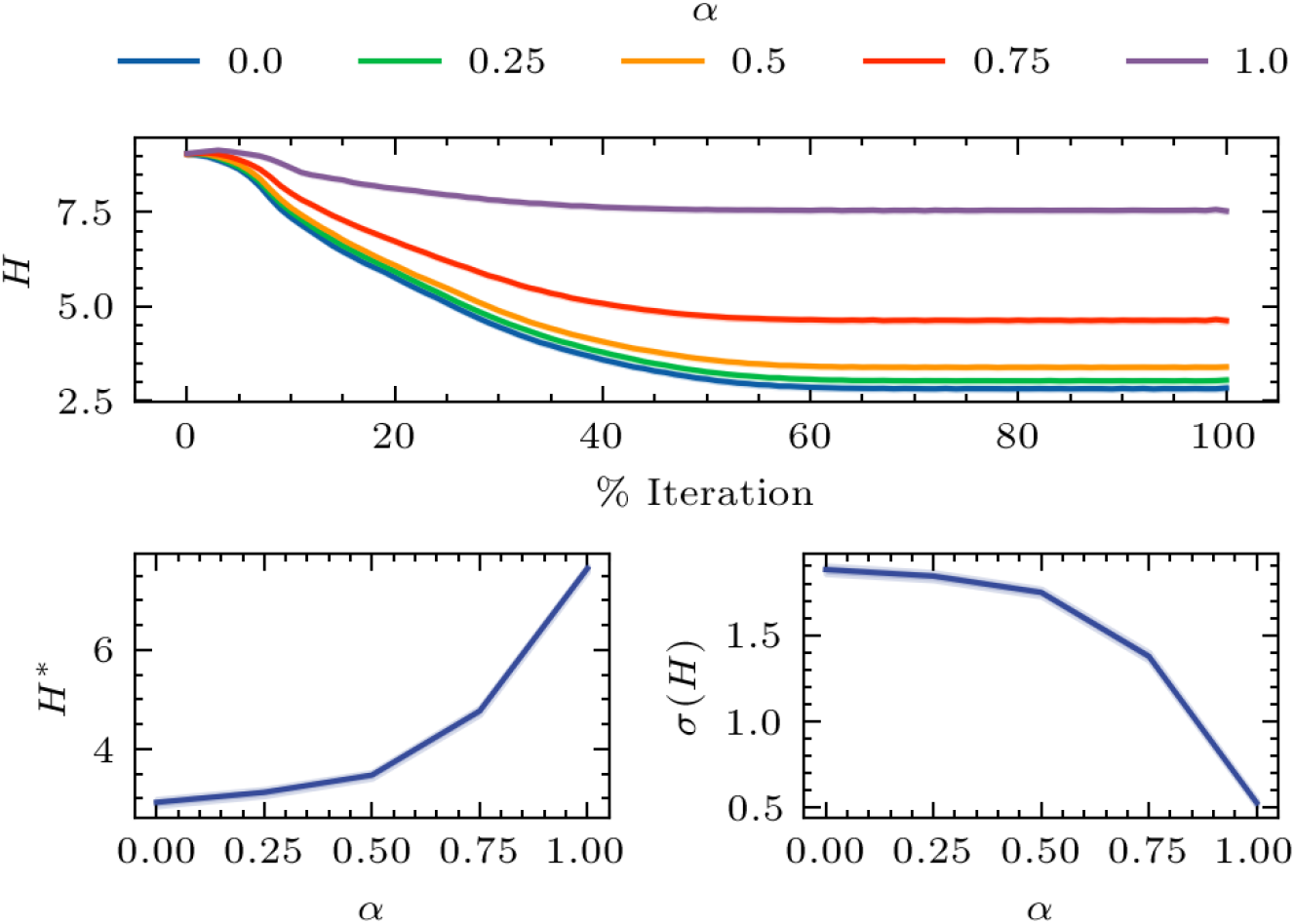
Line plots for the Shannon Entropy (*H*) per % iteration, converged *H* (*H*^*^), and standard deviation of *H* (*σ*(*H*)) during training for structures of length *N* = 6.

We expand upon these results by evaluating inferred contact probabilities from the sampled pairwise distances. Structures that contain any pairwise distance 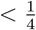 (spatial clash) are ignored. The inferred contact probability given the pairwise distance *d*[*i, j*] was motivated by Ref. ^27^ and is defined as the following:

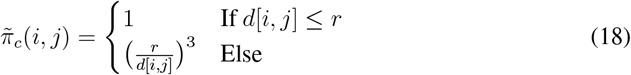

We evaluate the inferred contact probabilities 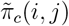 by calculating the average precision^106^ (AP) and area under the curve^106^ (AUC). We set the average inferred contact probabilities of the positive labeled contacts and negative labeled contacts as the the positive and negative predictions, respectively, hence making the labels perfectly balanced. We also computed the maximum and mean of the average inferred probabilities for positive labeled contacts. All four metrics are displayed in Figure 9 for targets with lengths *N* ∈ [6, 11).

**Figure 9:**
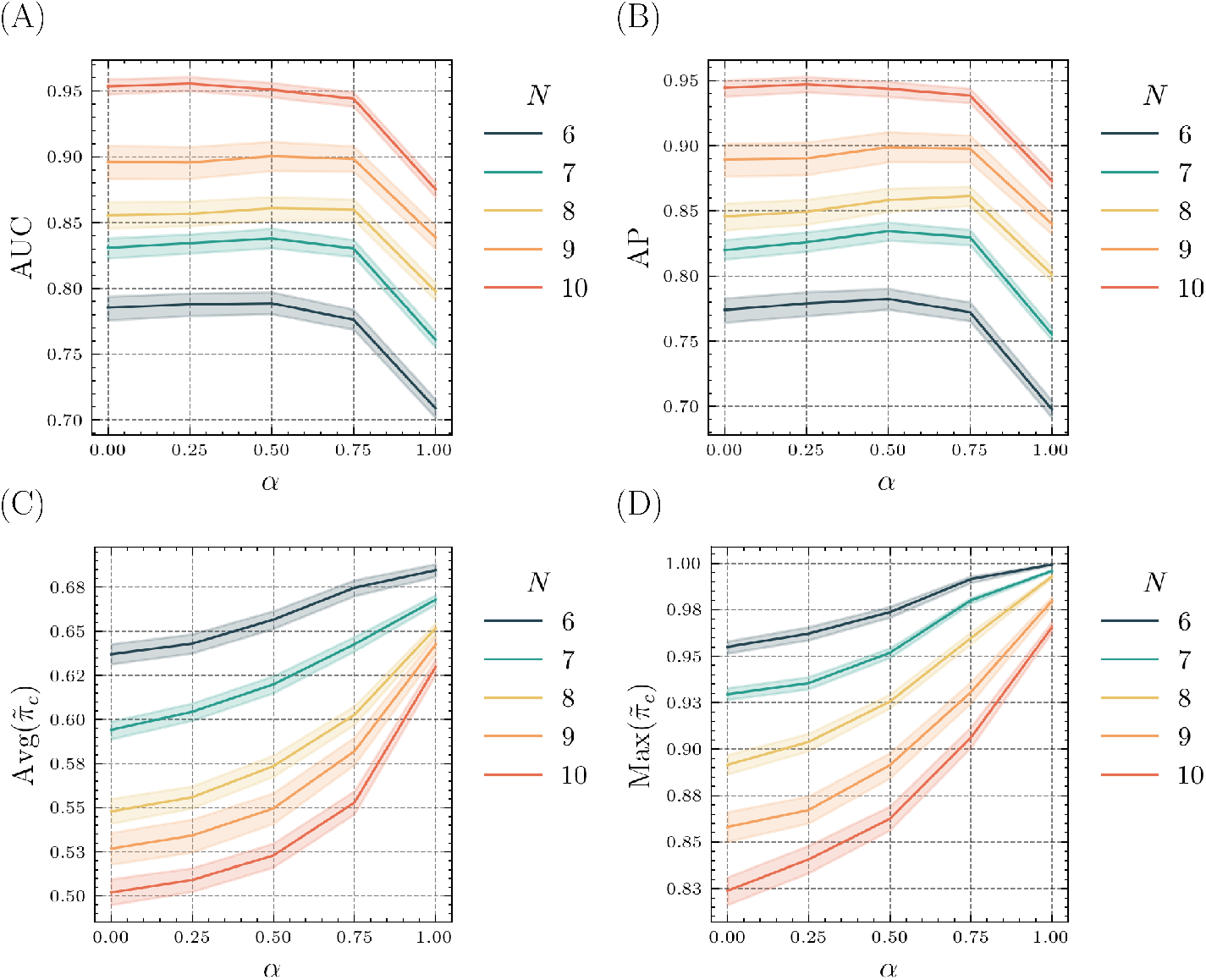
Line plot and confidence interval of the area under the curve (A), average precision (B), average (C), and maximum (D) inferred contact probabilities.

What we observe supports the relation of *α* to the single-cell and bulk Hi-C experiments. When *α* is smaller (single-cell) more precise inferred contacts are recovered. When modeling a bulk Hi-C experiment, more variations not only in the conformations is expected (single-cell), but also in the recovered contacts, as the contacts are captured from a large heterogeneous set of single-cell conformations^107^ (Bulk Hi-C). In other words, we expect and observe the precision and ranking metrics (Subfigures 9.A and 9.B) to be stronger for lower values of *α* (single-cell) and the recall metrics (Subfigures 9.C and 9.D) to be stronger for higher values of *α* (Bulk Hi-C).

We also define two variants of the Dice-Sørensen Index (DSI)^108,109^, weighted (*DSI*_*w*_) and maximum-threshold (*DSI*_*m*_), see Equations 19 and 20. Both assess the contact-recovery performance, with higher values indicating a stronger ability to recover target contacts.

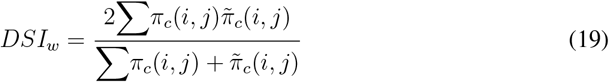

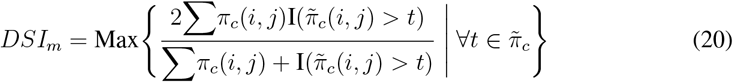

From Figure 10, it is clear that the aggregation parameter *α* can cause changes in the performance metrics, where both *α* = 0 and *α* = 1 are not optimal. This suggests that the ability to model and fine-tune the amount of aggregation can offer notable advantages, depending on the use-case.

**Figure 10:**
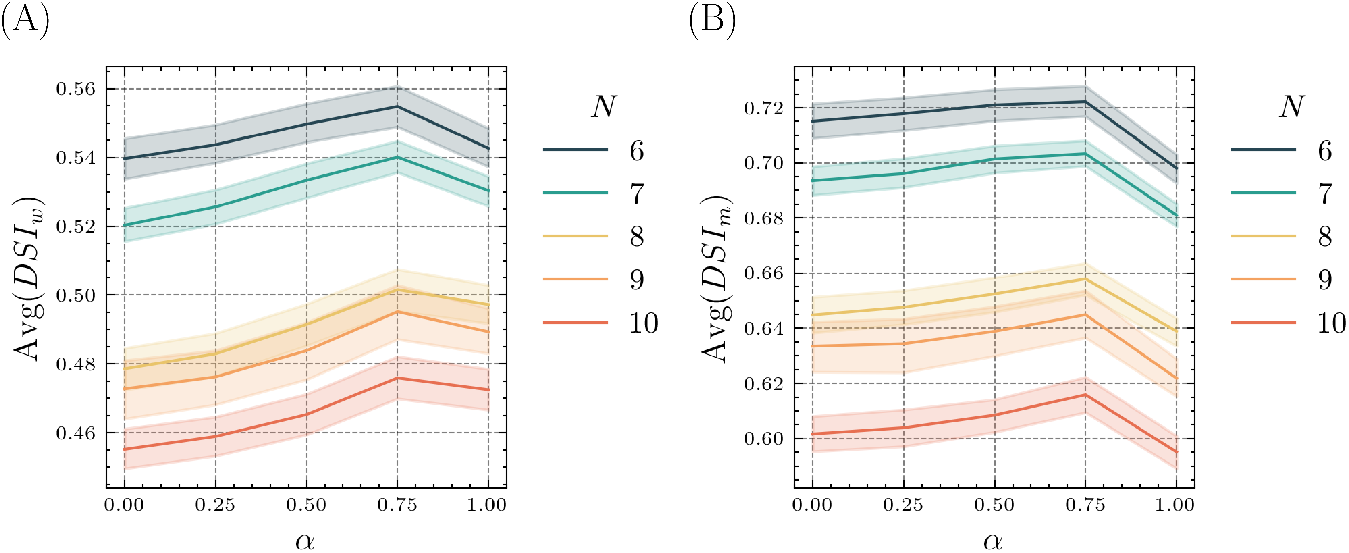
Line plot and confidence interval of the weighted (A) and maximum threshold (B) Dice-Sørensen index^108,109^.

#### 3.1.2 Landscape analysis of the aggregation parameter *α*

To gain further insights into the affect of modeling aggregation, we performed exploratory landscape analysis (ELA) and principal component analysis (PCA)^110^ over ℒ (*θ, α*). Our findings suggest ℒ (*θ, α*) becomes less rugged as we increase *α*.

Following the procedure described in Ref.^88^, we computed the empirical maximum information content (*H*_*M*_) over the parameter space. The number of parameters, *p*, was used to determine the number of samples, 128*p*. Similarly, we computed 64 random walks each with a length of 64*p*. We notice that *H*_*M*_ tends to decrease as we increase *α*, see Figure 11. The value of *H*_*M*_ is related to the ruggedness of the landscape^88,111,112^, with higher values suggesting a more rugged landscape, and hence are generally more trainable.

**Figure 11:**
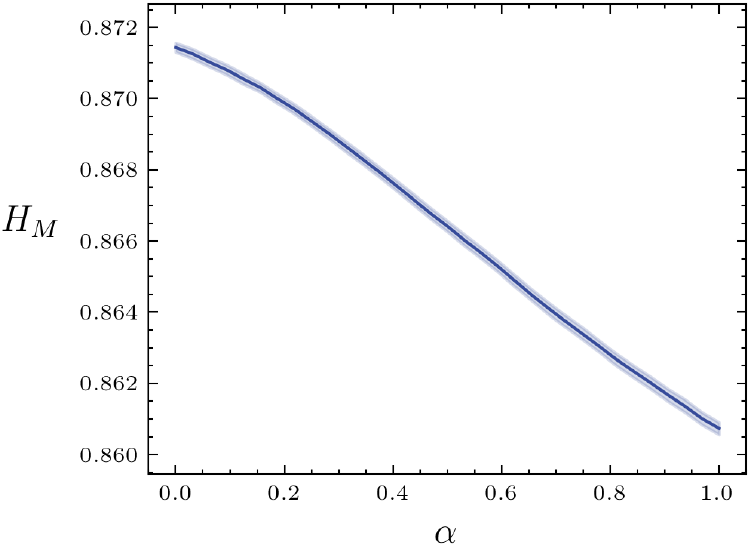
Confidence interval and line plot of empirical *H*_*M*_ values across a random sample of groups of length 6 structures.

We further observe this visually through PCA embeddings of the objective function landscape, see Figure 12.

**Figure 12:**
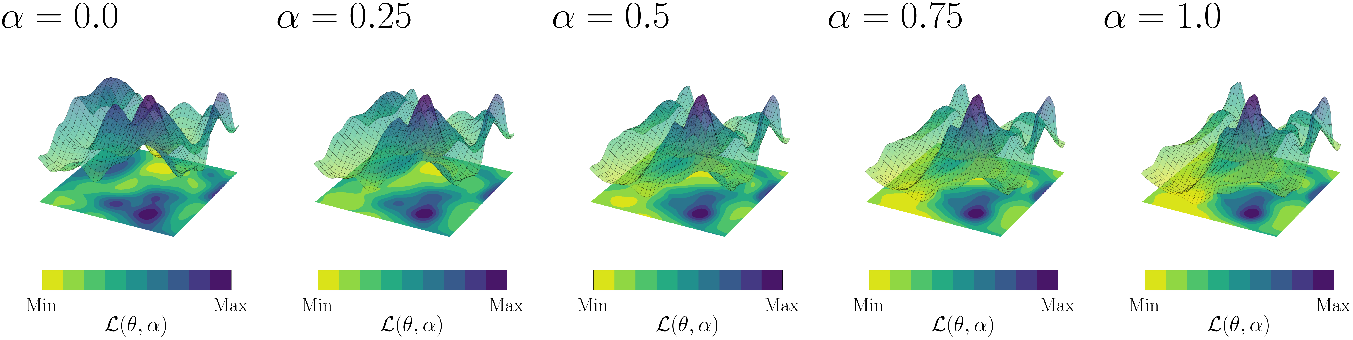
PCA embeddings of the learned parameter landscapes with *α* = 0 for a single target group of length *N* = 6 structures. The PCA embeddings are fixed, with the Z-axis denoting the loss function ℒ (*θ, α*). The loss function for each landscape is calculated explicitly with different values of *α* to show the change in local minimums of ℒ (*θ*, 0) as *α* is increased. Each loss is independently normalized to the same scale (linearly) allowing for clear comparison.

As we increase the value of *α*, the landscape becomes more flat and evenly distributed around the local minimums. Generally, this would be viewed as an increase in optimization landscape difficulty. However, we observed in Section 3.1.1 that the proposed algorithm still behaves as expected for different values of *α*. This suggests that depending on the use-case and problem instance, the gained utility of increasing *α* could outweigh the slight increase in landscape difficulty.

### 3.2 Evaluation on experimentally-obtained genomic contact data

To further validate our methodology, we modeled the 3D structure of a low resolution bulk Hi-C matrix with the proposed VQA. Captured contact frequency and euclidean distance form an inverse power law relationship^28–30^:

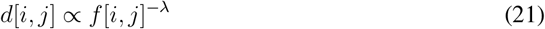

The optimal power *λ* > 0 can vary between datasets. Let us denote the minimum non-zero contact frequency detected between loci *i* and *i* + 1 as *f*_unit_. We define the contact matrix *π*_*c*_ in the following way:

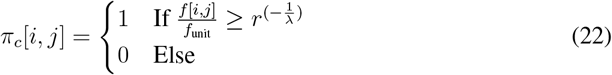

Similar to Equation 18, we define the inferred contact probability, 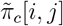,given the pairwise distance *d*[*i, j*] as the following:

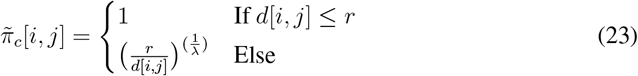

We used Cooler^113^ to produce a low-resolution (10MB) balanced contact matrix of chromosome 18 for the GM12878 cell-line^9^, see Table 1.

**Table 1:** Genomic data configurations for the GM12878 cell-line.

| Chromosome | Resolution | Polymer Length | $\lambda$ | #Contacts |
| --- | --- | --- | --- | --- |
| 18 | 10MB | 8 | 1/3 | 17 |
| 18 | 10MB | 8 | 1/2 | 11 |
| 18 | 10MB | 8 | 2/3 | 8 |

For every configuration defined in Table 1 we ran ten trials of our algorithm with noise-induced simulators and a noise-free simulator, as in Section 3.1.1. We then prepared and sampled the optimal pre-trained parameters, for each of the ten trials, to see if the classically learned parameters can effectively replicate the distribution of learned genomic conformations on a real quantum device. We ran our experiments on the physical *ibm_sherbrooke*^94^ quantum device, of which the noise was modeled during simulations. However, due to high computational resource utilization for noisy-simulation and long wait times for access to the real quantum device, we only performed experiments with *α* = 0 and *α* = 1.

We first investigate the empirical Shannon Entropy 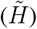 for the set of sampled states 𝕊^*^, see Equation 24 and Figure 13. We observe the same patterns in Figure 13 as in Figure 8.

**Figure 13:**
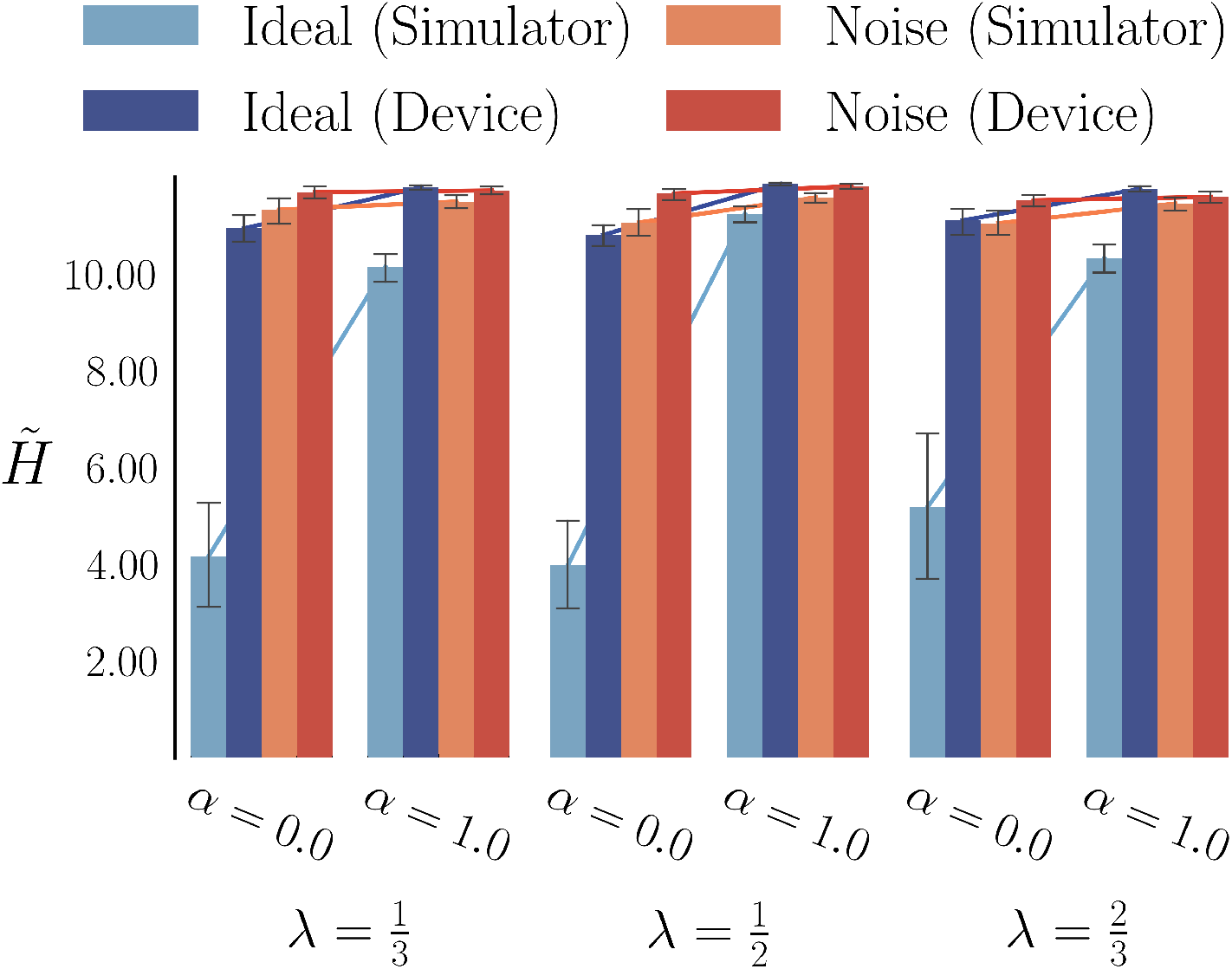
Bar plot with confidence interval of the empirical entropy 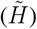.

However, it is clear that the empirical entropy from the noisy simulator and real device are higher than that of the ideal parameters on the noise-free simulator. This is expected, as noise naturally increases the uncertainty of the distribution, and hence increases the entropy. This suggests that in general noise impacts the learned distribution and thus more precise devices can improve the utility of the samples.

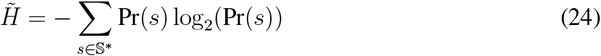

In Figure 14, we plot the AUC, AP, and recall metrics for the samples from the optimal parameters on the real device and learned parameters on the simulators (ideal and noisy). The metrics were computed as in Figure 9.

**Figure 14:**
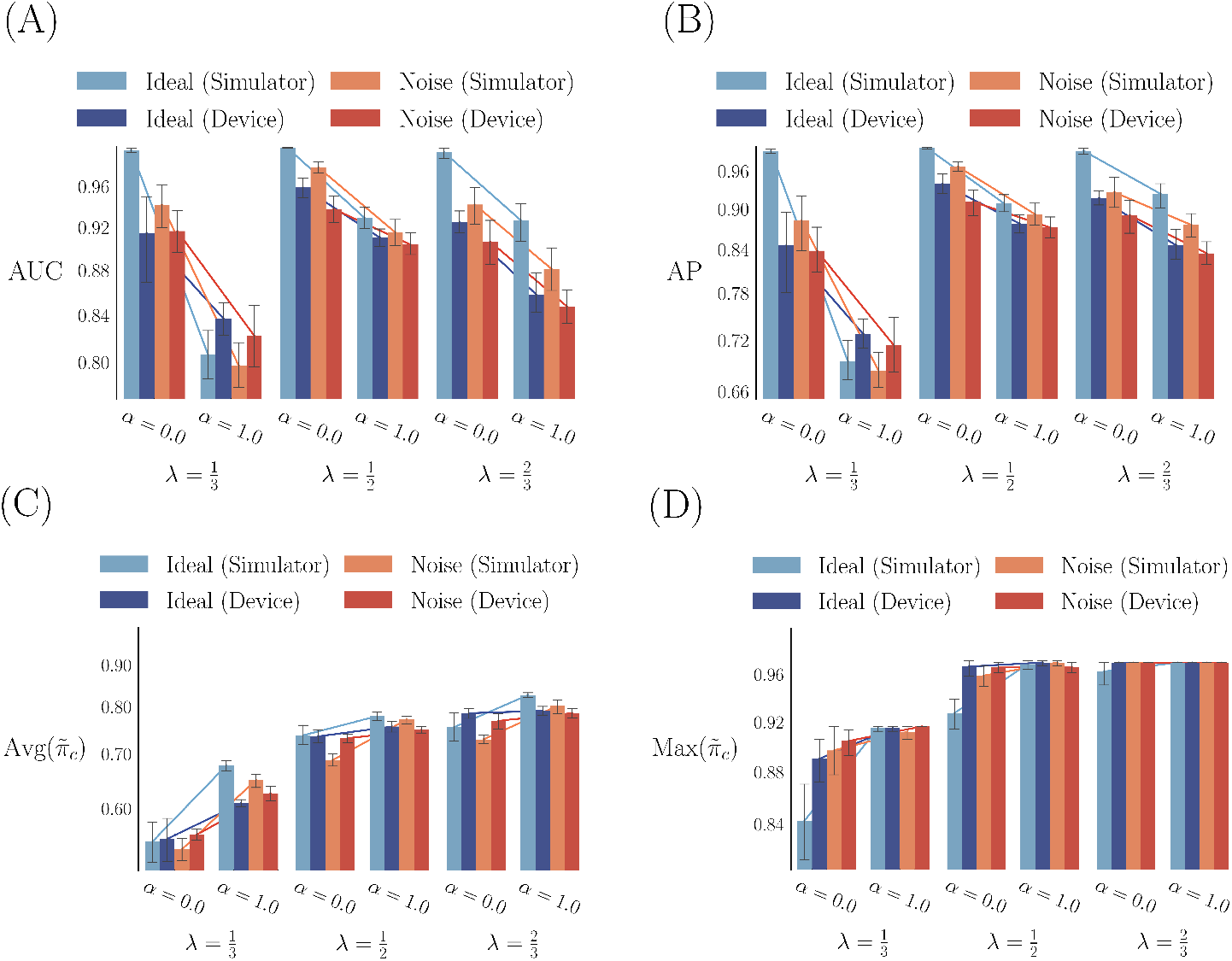
Bar plot with confidence interval of the area under the curve (A), average precision (B), average (C), and maximum (D) inferred contact probabilities. Parameters (*θ*) were found through Qiskit simulations^94^, with the optimal parameters (*θ*^*^) run on both the simulators and the *ibm_sherbrooke*^94^ quantum device.

Similar to the empirical entropy analysis, we also observe the same patters in Figure 14 as in Figure 9. Interestingly, while noise can affect the entropy of the distribution significantly, it has less of an affect on the expected trends for AUC, AP, and recall measures. This demonstrates that even in the presence of noise, the learned parameters still can resemble the distribution of conformations for single-cell (*α* = 0) and bulk (*α* = 1) Hi-C experiments. While it is still necessary to conduct larger scale experiments in the future, given access to and improvements in quantum hardware, these are promising and encouraging results.

## 4 Discussion

In this paper, we propose a novel variational quantum algorithm for reconstructing 3D genome structures from experimental genomic contact data. Our extensive analysis supports the notion that our algorithm can effectively model and sample from the distribution of genomic conformations. We have also shown that our methodology can theoretically model the conformational space expected for both single-cell and bulk experiments.

There are several potential benefits for reconstructing 3D genome structures with a variational quantum algorithm, particularly when modeling conformational ensembles. When training the proposed VQA, the number of parameters grows linearly with respect to the number of genomic loci, which is only slightly larger than that needed to store a few genomic conformations classically. This suggests that when modeling bulk Hi-C data, we can de-convolve genomic structures into an ensemble without having the optimization memory requirements (parameters) grow with respect to the size of the ensemble. This same idea applies to single-cell Hi-C, where we embed a likely ensemble of genomic structures corresponding to a single-cell Hi-C matrix. In other words, by using parameterized quantum circuits, we embed and optimize over the space of conformational ensembles without requiring a significant increase in parameters. Furthermore, sampling from a quantum machine is theoretically very efficient and thus could offer benefits compared to parallel classical sampling (simulated annealing).

Another potential utility of the proposed algorithm is the ability to replicate and fine-tune the learned distribution of genomic conformations. There is evidence for the relation between epigenetics and 3D genome conformations^7,114,115^. For example, it is well established that CTCF binding sites (detected by ChIP-Seq^116^) are related to TAD boundaries, which have a precise mathematical connection to captured genomic contacts^117^. Another example is chromatin accessibility profiling^118^, which provides insight into the degree of chromatin packing (radius of gyration), i.e., euchromatin and heterochromatin. Furthermore, epigenetic data has been used to predict chromosomal contacts and vice versa^27,119–122^, supporting their connection. We can enhance our objective function by directly evaluating inferred structures against epigenetic data by the related three-dimensional properties of the genome. Thus, by preparing |*ψ*^*^⟩ (assuming reasonable noise levels), we can explore a variety of biological phenomena (fine-tune) without being limited to a single reference structure. However, a larger, higher-resolution polymer than those experimented with in this study would be needed to model these properties. Hence, classical simulation (simulating quantum circuits on a classical computer) and current quantum hardware cannot support experimenting with these aforementioned properties at this moment. With forthcoming advancements in quantum hardware, future models can further explore and model the relationship between the 3D genome and other biological processes with high-throughput epigenetic data.

The results from this study serve as a starting point for highlighting the conceivable impact quantum computation can have on three-dimensional genomics. We expect future advancements in quantum technology to come with much promise for insights into the 3D genome and its respective regulatory landscape.

## 5 Acknowledgements

This work was supported by the National Institute of General Medical Sciences under Award Number 1R35GM137974 to Z.W. The physical quantum devices utilized in this study were accessed via the open plan provided by IBM’s Quantum Platform.

## 6 Author contributions

A.J.S and Z.W. conceptualized and developed the methodology. A.J.S implemented and wrote the code. A.J.S prepared and visualized the data and results, respectively. A.J.S wrote the original draft of the manuscript. Z.W. and A.J.S revised and edited the manuscript. Z.W. supervised and provided funding. All authors read and approved the final manuscript.

